# Signed motif analysis of the *Caenorhabditis elegans* neuronal network reveals positive feedforward and negative feedback loops

**DOI:** 10.1101/2025.01.09.632090

**Authors:** Gabor S. Szilagyi, Attila Gulyas, Zsolt Vassy, Peter Csermely, Bank G. Fenyves

## Abstract

**Background:** Nervous systems are complex biological networks with largely unknown structural and functional characteristics. Motif analysis is a robust tool that can reveal the unique aspects of connectivity in a complex network. An ideal candidate for motif analysis is the connectome of the nematode *Caenorhabditis elegans*, which is the first fully reconstructed nervous system.

**Results:** We performed, for the first time, edge-polarity based signed motif analysis on the *C. elegans* connectome using recent data on the connection signs of this network and a novel structure-preserving randomization method. We identified 56 significantly over– and 1 underrepresented three-node signed motifs and revealed that certain motifs (e.g., positive feedforward, negative feedback, disinhibitory feedback, and incoherent feedforward loops) are overabundant in the *C. elegans* connectome. We further distinguished nodes by their corresponding neuron modalities (e.g., sensory vs. motor neurons), and found that each significant feedforward and feedback loop has a characteristic neuronal layout.

**Conclusions:** Our findings demonstrate the importance and potential of signed motif analysis in understanding biological networks. The motif enumeration tool and definition system we developed can be used to analyze signed motifs in other complex networks.

## Background

Nervous systems are complex biological networks that generate behavior through neural activity mediated by synaptic connectivity. Biological networks share common global (e.g., small-worldness, cost-efficient wiring) and local (e.g., clustering, modularization) properties that allow optimal information processing [1–4]. Importantly, detailed structural characteristics can be revealed by investigating a network’s composition from smaller building blocks, called motifs (Fig. 1A). Motifs are frequently occurring subgraphs that correspond to different biological functions [3]. A subgraph is a graph whose nodes and edges are subsets of another graph, with two main types: induced and partial [5]. A subgraph is induced when it consists of a subset of nodes from a network and all the edges that connect them. On the other hand, partial subgraphs contain only some of the edges connecting the chosen nodes (Fig. 1B). The classical concept is that if a specific subgraph occurs in a network more frequently than expected, that subgraph might play a crucial role in the network; hence, it is called a motif. Motifs have been examined in a variety of real-world and, more importantly, brain networks [3, 4, 6]. Like the networks themselves, motifs are characterized primarily by their nodes and edges.

**Fig. 1.**
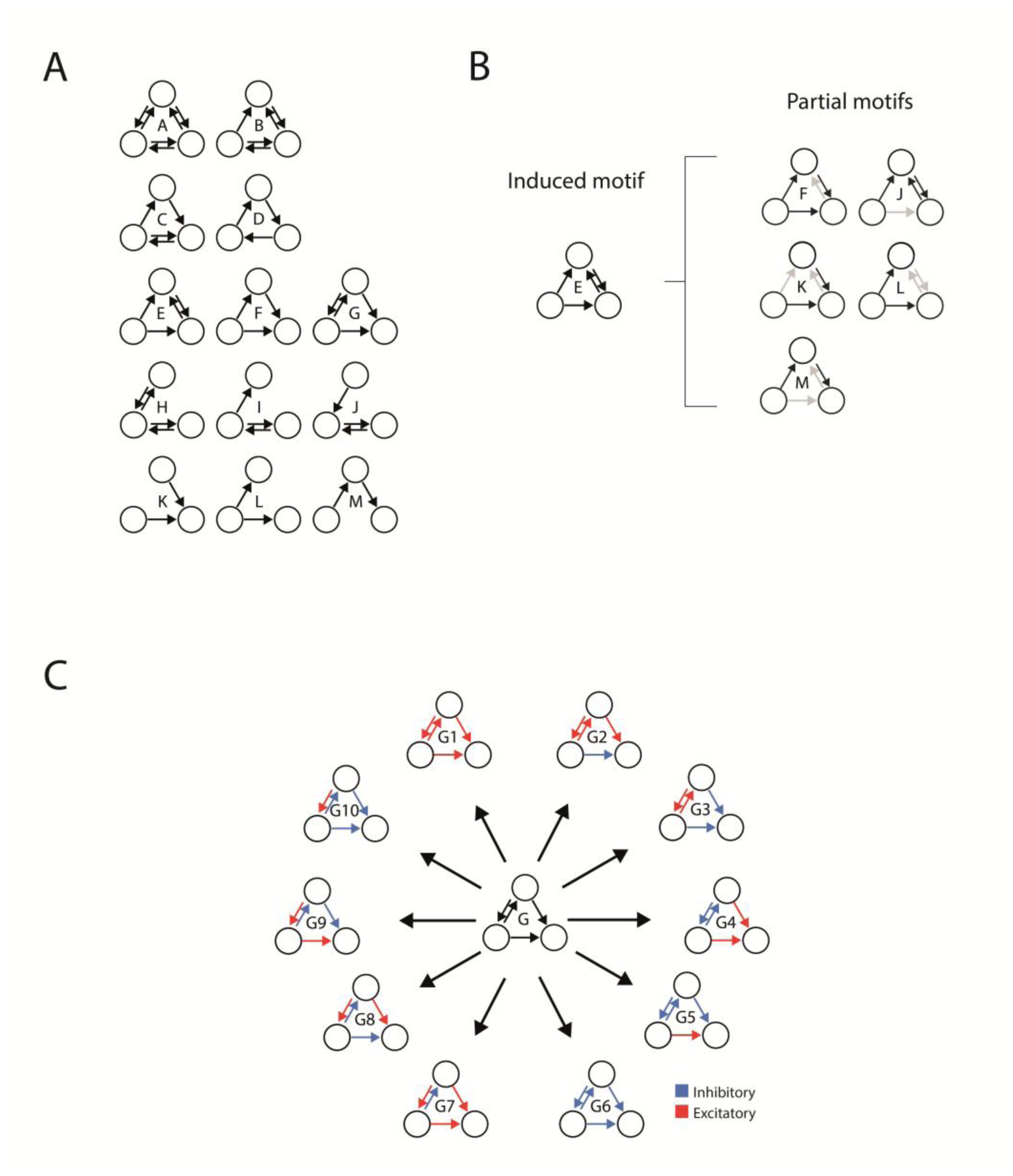
Overview of motif definitions. A) Three-node motif structures with directed edges. B) Induced and partial motifs. Partial motifs are subgraphs of their corresponding induced motif. For example, the induced motif E contains five different partial motifs. A similar concept is discussed in McDonnell et al. (2014) and Sporns & Kötter (2004). C) Definition of signed motifs by coloring the edges (of motif G in the example). An edge can be either excitatory or inhibitory.

Particularly in brain networks, edges can be labelled (colored) by the polarity of the connection they represent, i.e., excitatory or inhibitory (Fig. 1C). This is important, as structurally identical motifs with different polarity patterns can have completely different biological functions. However, edge polarity-labeled (i.e., signed) motif analysis is rarely performed on neuronal networks, since large-scale polarity data are lacking in most species.

Even though there are now multiple known connectomes [7, 8], the neuronal network of the nematode *Caenorhabditis elegans* was the first completely reconstructed cell-level representation of a living organism’s nervous system [9, 10], consisting of 302 neurons and ∼5000 connections in a hermaphrodite. In previous work on truncated and partial worm connectomes, some three– and four-node network motifs were found to be overrepresented [3, 10–14]. At the same time, changes in motif occurrences during the animal’s development have also been observed [13]. Motif analysis has revealed the backbone structure of the connectome [15]. Some further functional analyses have been reported in recent studies, which have colored the nodes by neuronal function and neurotransmitter expression or labeled the edges by synapse type [16–18].

In a recent work, we published a comprehensive dataset of synaptic polarities for the *C. elegans* chemical synapse connectome [19]. Using presynaptic neurotransmitter and postsynaptic receptor expression data, we predicted the polarities (excitatory, inhibitory, or complex) of more than 15,000 chemical synapses.

In this study, we analyzed the distribution of signed motifs in the *C. elegans* connectome using previously published connectome-scale synaptic polarity data. We developed a novel method and a computational tool for signed motif analysis and applied them to the edge polarity-based signed chemical synapse neuronal network of *C. elegans.* We carried out structure-preserving network randomization to generate null models for motif analysis. We showed that some signed three-node motifs (most notably, negative feedback and positive feed-forward loops) are significantly overrepresented in the chemical synapse subnetwork of the *C. elegans* connectome. To assess robustness, we performed sensitivity analyses to evaluate the effect of connections with unknown or potentially falsely predicted polarities. 75-80% of the motifs that were found to be overrepresented in the network were robust during these validity tests, while the only notable motif lost during this was the coherent positive feedforward loop. Our study identified multiple statistically overrepresented polarized motifs without a known function in the *C. elegans* connectome, whose biological relevance remains to be determined experimentally in future studies.

## Methods

### Description of the *C. elegans* connectome data

The WormWiring connectome reconstruction (http://wormwiring.org) of the adult hermaphrodite worm comprises 3,638 chemical connections (20,589 synapses) among 297 neurons (CANL, CANR, PLML, PLMR, and M5 neurons are not connected by chemical connections and were therefore excluded from the analysis). We used the previously published signed neuronal network, in which synaptic signs were predicted from cellular-level data on neurotransmitter and receptor expression [19]. In the original work, synapses were predicted to be excitatory or inhibitory when only cation– or anion-channel receptor genes matched the presynaptic neurotransmitter, respectively. Synapses were labelled as complex if both types of receptor genes were expressed on the postsynaptic cell. Finally, the synapse remained unpredictable if no receptor gene matched the presynaptic neurotransmitter. In the present work, complex synapses were relabeled as excitatory, inhibitory, or unknown, based on the ratio of anion– and cation-gated postsynaptic receptor expression of the given neurotransmitter (e.g., connections that had excitatory>inhibitory postsynaptic receptors were defined as excitatory) to reduce combinatorial complexity, similarly to a method used previously [20]. If the number of excitatory and inhibitory receptors were the same, the synapses were relabelled as being unknown. Hence, in this work, a network edge was colored with one of three labels (Additional file 1). We extracted neuron modalities (i.e., sensory, inter, motor, or polymodal) from Wormatlas (http://wormatlas.org). Neurons with more than one functional modality (e.g., inter– and motor) were labeled polymodal.

### Definition and identification of signed three-node motifs

A definition system for directed, edge-colored motifs with three nodes was established. Edges were signed as excitatory, inhibitory, or unknown, resulting in 710 motifs (Additional file 2). Rotational and reflectional symmetries were considered as the same motif. Motif search was performed using a custom-developed algorithm [21] that seeks induced subgraphs, identifies all three-node, edge-colored motifs in the target network, and categorizes them as one of the 710 motif types. Motif counts and exact motif locations are described in Additional file 3. The results of the induced subgraph analysis are presented in Additional file 4. Our algorithm was further used to convert induced motif counts to partial motif counts (Additional file 5).

For further analysis of feedforward and feedback loops, we assigned a modality to each neuron based on the data available (http://wormatlas.org). If a neuron exhibited multiple different modalities (i.e., can be both a sensory neuron and an interneuron), it was labelled “Polymodal”. To reduce the number of modality layouts, if a motif contained at least one polymodal neuron (i.e., a neuron without a specific modality), it was labeled “P” without further differentiation.

### Randomization methods and statistical analysis of three-node motifs

Although the conceptual framework of motif analysis and null model generation in non-signed neuronal and non-neuronal networks is well established [3, 10, 22], the approach to edge-colored (signed) motif analysis is less straightforward. Therefore, we developed a conceptual framework for our study. The hypothesis of our null model was that synaptic polarity is distributed independently of the connectome’s structural wiring. To establish this biologically meaningful null model, we decided to keep the network wiring (i.e. physical structure) of the connectome constant, and to perturbate edge polarities indirectly via randomizing neurotransmitter expression (a main determinant of synaptic polarity [19]) of all neurons and then re-deriving synaptic polarities from the randomized neurotransmitter–receptor pairings (Fig. 2A). This approach preserved (i) the network topology, (ii) the relative abundance of each neurotransmitter across the network, and (iii) the postsynaptic receptor-expression profiles. This logic is consistent with the standard framework for null models in network neuroscience, in which specific architectural properties are selectively preserved while others are systematically randomized [22]. It is also the logic of colored motif analysis, in which the null is constructed so that node or edge labels are independent of topology, and rejection of the null indicates that the labels and the topology are organized non-independently [23]. Notably, this method yields a varying ratio of excitatory to inhibitory connections in random networks compared with the original connectome (Fig. 3). Nevertheless, compared with simpler randomization methods (such as shuffling edge polarities directly – also reported in this study, see “alternative randomization”), this method yields random networks with biologically plausible underlying patterns of neurotransmitter ratio and receptor expression. Since the connectome’s structural hardwiring was preserved, absolute motif counts could be used for analysis. A total of 1,000 random networks were generated and used as a null model (Additional file 1).

**Fig. 2.**
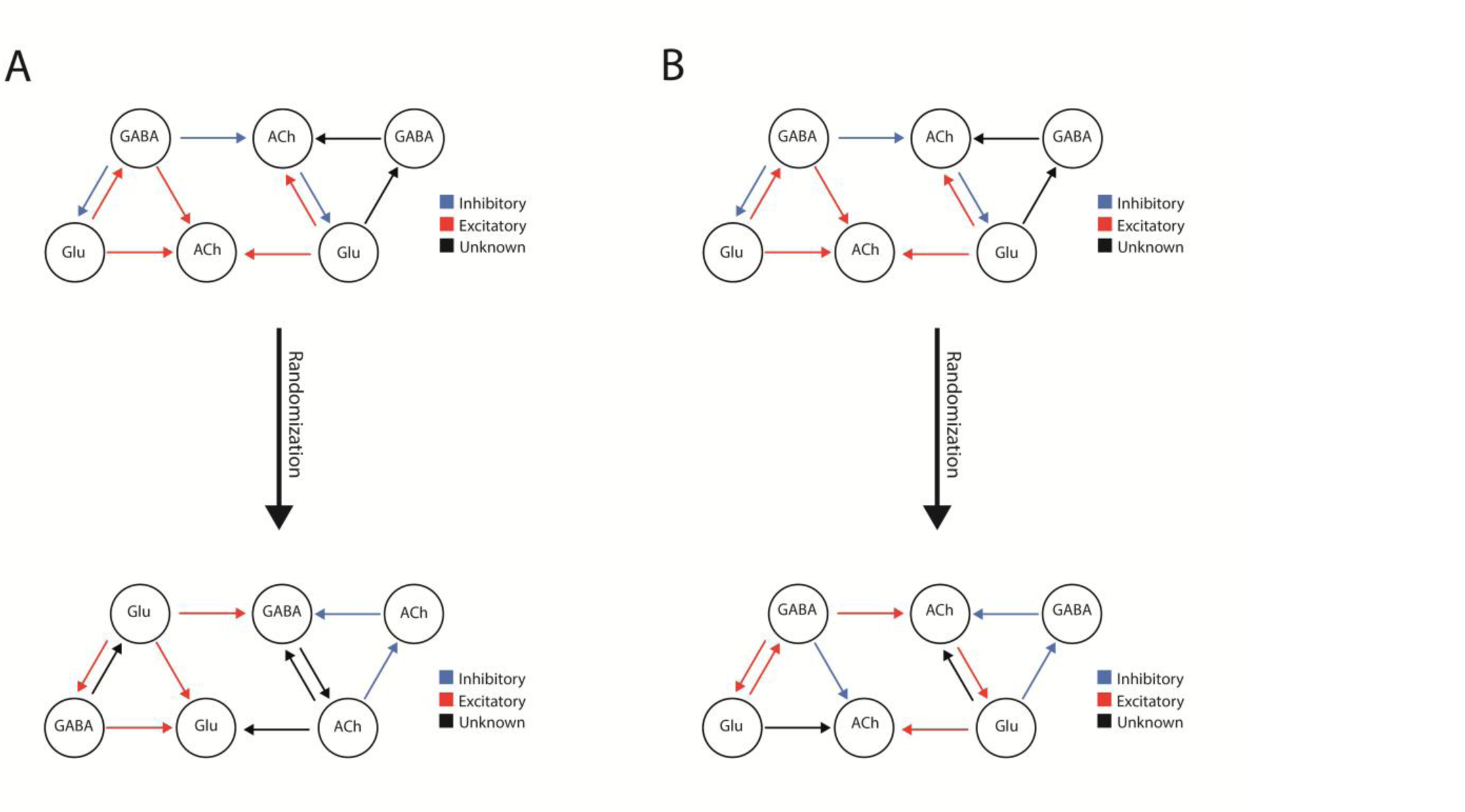
Neurotransmitter– and edge polarity-shuffling randomization methods. A) Neurotransmitter-shuffling randomization. In this method, the neuronal neurotransmitter expression patterns were randomized. Because receptor expression patterns were kept constant, the resulting networks differed in the numbers of excitatory, inhibitory, and unknown edges. B) Edge polarity-shuffling randomization. In this method, excitatory, inhibitory, and unknown synapses were randomly assigned across the network. This resulted in a null model that preserved the original network’s excitatory-to-inhibitory ratio. The red and blue arrows represent excitatory and inhibitory synapses, respectively, while black arrows stand for connections with unknown polarity.

**Fig. 3:**
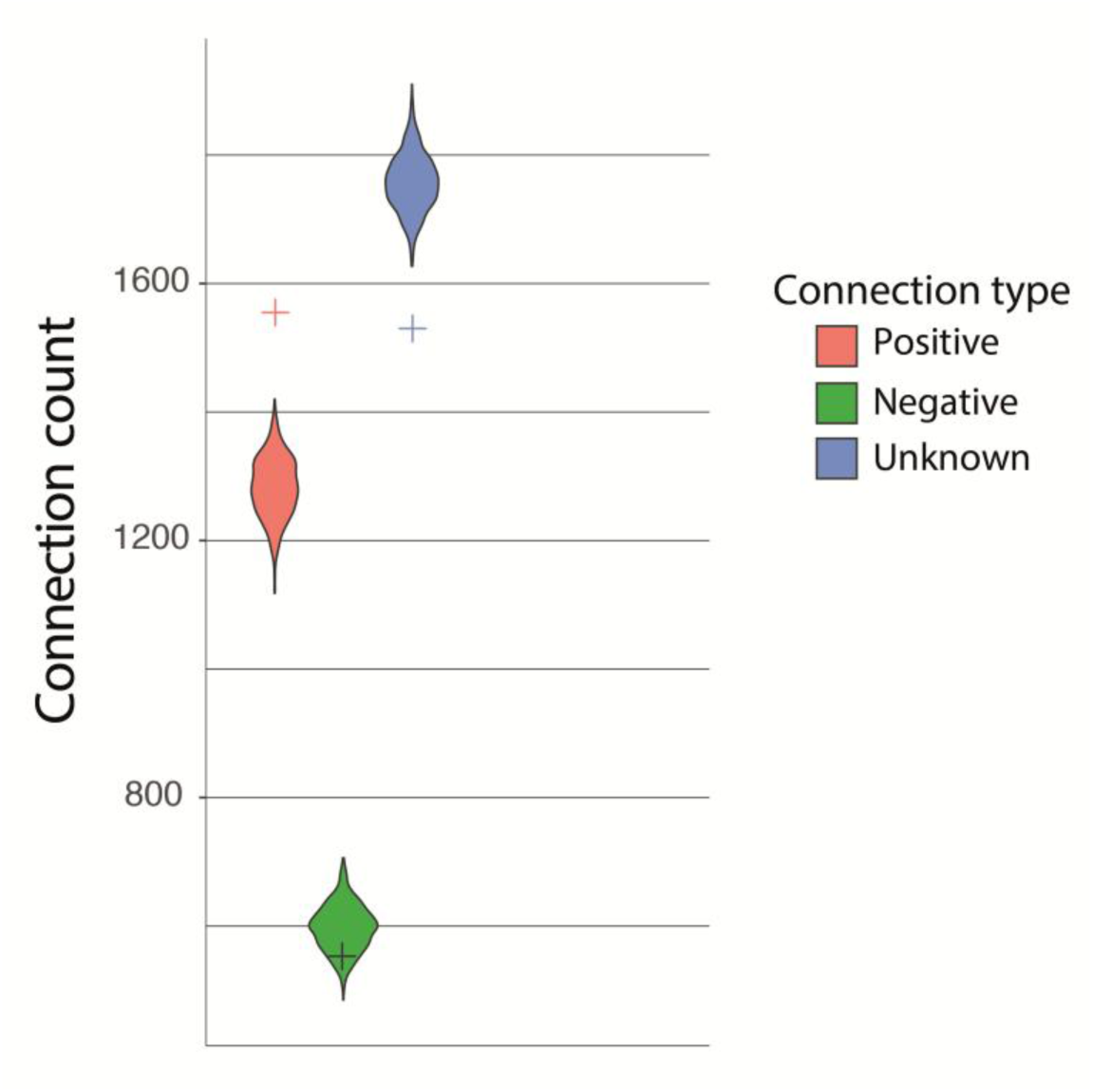
Distribution of different edge polarities among the connectome (crosses) and the generated 1,000 random networks (violin plots). Random networks were generated by randomizing the neurotransmitter expression of neurons in the connectome; therefore, only edge polarities changed, and the numbers of edges and nodes remained the same.

Even though our randomization method follows the biological nature of the connectome, randomizing neurotransmitter expression increased the number of connections with unknown polarity due to potential receptor mismatches. Moreover, the network’s excitatory-to-inhibitory ratio varied widely across the randomized networks. To address this, we generated an alternative null model by simply randomizing the edge polarities (a widely used standard in network analysis) (Fig. 2B, Additional file 6). This resulted in 1,000 random networks with the same number of excitatory and inhibitory connections as the connectome. However, this results in greater variability among the null-model networks, since (contrary to the primary randomization method) neurons expressing only anion– or cation-gated receptor channels can receive synaptic input of the opposite polarity than their receptor pattern suggests.

In the formal analysis, we tested an alternative hypothesis where a colored motif in the *C. elegans* neuronal network is significantly over– or underrepresented compared with the null model, as established previously [18]. As an established method to assess the statistical significance of individual motifs [3, 24], for each motif, the Z-score was calculated using the following formula:

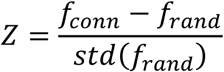

where *f_conn_* is the frequency of the motif in the connectome, *f_rand_* is the average frequency of the motif in the random networks, and *std*(*f_rand_*) represents the standard deviation of the motif’s occurrence in random networks.

After calculating p-values from the Z-scores, we performed false discovery rate correction using the Benjamini-Hochberg procedure. Motifs with a corrected p-value under 0.05 were considered significantly over– or underrepresented (95% confidence interval, alpha=0.05).

### Analysis of two-node motifs

For two-node motifs, we colored the edges by synaptic polarity and the nodes by neuronal modality. The list of neuronal modalities is provided in Additional file 7. All nodes were labeled using one of the four possible neuron modalities (“sensory neuron”, “interneuron”, “motor neuron”, or “polymodal neuron”) based on their role in the connectome (http://wormatlas.org). Partial and induced motif analyses were performed as described for three-node motifs, using the same null model for statistical analysis. Results are presented in Additional file 8. All tools used are available at [21].

### Validation

We performed two sensitivity analyses to assess the impact of our major limitations on the study results. In the first analysis, we eliminated unknown connections by randomly assigning them an excitatory or inhibitory polarity while maintaining the excitatory/inhibitory ratio of the network. Using this approach, we generated 1,000 networks that we used as a new null model to examine changes in both induced and partial motif occurrences relative to the original network (Additional file 9). Because this method necessarily increases the number of motifs without an unknown edge, we used motif ratios rather than absolute counts for statistical analysis.

In the second analysis, we aimed to evaluate the effect of using predicted polarities that may be inaccurate. To address this, we randomly changed the polarity of edges in the connectome at a 10% noise level. Using this method, we generated 1,000 networks for induced and partial motif enumeration and then statistically compared motif counts with those of the connectome (Additional file 10). In both analyses, Z-scores were calculated, and the Benjamini–Hochberg procedure was applied to control the false discovery rate.

### Software

The motif search algorithm was developed in Python (3.10.11). Randomization of the network and polarity prediction was performed in R (4.2.2). Statistical tests were performed in R (4.2.2) and Microsoft Excel (16.84). All the scripts are available at public repositories [21].

## Results

### Induced subgraph analysis

The chemical synapse connectome of *C. elegans* consists of 3,638 connections, of which 1,555, 553, and 1,530 were labeled excitatory, inhibitory, or unknown, respectively, based on our previous work [19]. The process is described in the Methods section (Additional file 1). In this network, we identified 64,962 individual three-node subgraphs (Additional file 3). Their wiring structure and edge labeling categorized these subgraphs into one of 710 unique motif types (Additional file 2). To establish a null model, we generated a set of 1,000 networks by randomizing the neurotransmitter expression of the neurons. This resulted in a varying ratio of excitatory and inhibitory connections among the random networks (Fig. 3), but the total number and structure of edges remained the same across all networks. In the randomized networks, the number of unknown connections increased due to shuffled neurotransmitter expression among neurons. This disruption causes some postsynaptic cells to receive insufficient input from presynaptic partners. Since more neurons have cation-channel receptors for exactly one neurotransmitter than neurons with anion-channel receptors for one neurotransmitter, this mismatch is more likely to affect excitatory synapses. Therefore, this set of random networks had, on average, fewer excitatory connections and more unknown connections than did the original *C. elegans* connectome (Fig. 3). We counted the occurrences of these motifs in the connectome (Additional file 3) and in a set of random networks, and calculated Z-scores for each motif to determine over– or underrepresentation (Additional file 4). We identified 38 over-and 9 underrepresented motifs. Six motifs were absent both in the *C. elegans* connectome and in the null model; thus, no Z-score was calculated. A total of 184 motifs were only absent from the connectome. We found that 23 of the overrepresented motifs and 1 of the underrepresented motifs had no unknown edges (Fig. 4).

**Fig. 4:**
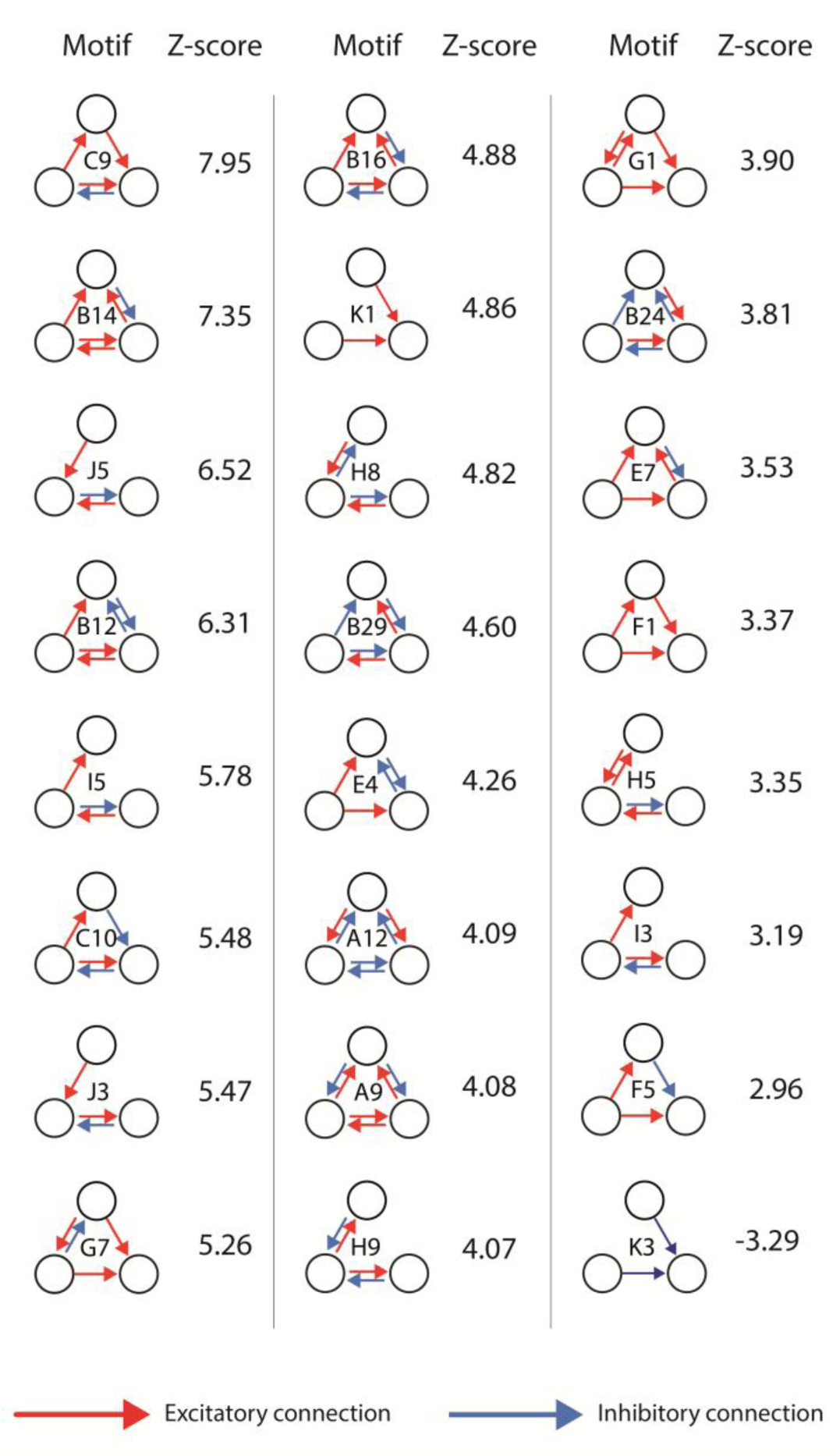
Induced motif analysis of the connectome. Using 1,000 random networks as a null model, induced motif analysis revealed 23 significantly overrepresented motifs and 1 underrepresented motif with no unknown edges. The red and blue arrows represent excitatory and inhibitory synapses, respectively. All data are available in Additional File 4.

### Partial subgraph analysis

Since smaller motifs can be embedded within more complex structures, partial subgraph analysis can provide a deeper, more functional understanding of information-processing networks [4, 5]. Therefore, we translated our findings of induced motifs to partial motif counts (Methods) and described 193,487 partial motifs in the connectome. We identified 49 significantly overrepresented and 8 underrepresented partial motifs (Fig. 5A, Additional file 5). Six motifs (motif A16, A33, A53, A83, A87, and A132) were absent both in the *C. elegans* connectome and in the null model; thus, no Z-score was calculated. Only one of them had no unknown edge (Motif A16). On the other hand, 153 motifs were absent only from the connectome, 22 of which had no unknown edge (Additional file 5). Forty percent of the identified partial motifs had only excitatory and inhibitory edges (Fig. 5B). Thirty-six of the significantly overrepresented motifs had no unknown edge, whereas all significantly underrepresented motifs contained at least one edge (Fig. 5C). Partial motifs with the highest Z-scores typically had one and only one inhibitory connection that provided a negative feedback function in the motif.

**Fig. 5.**
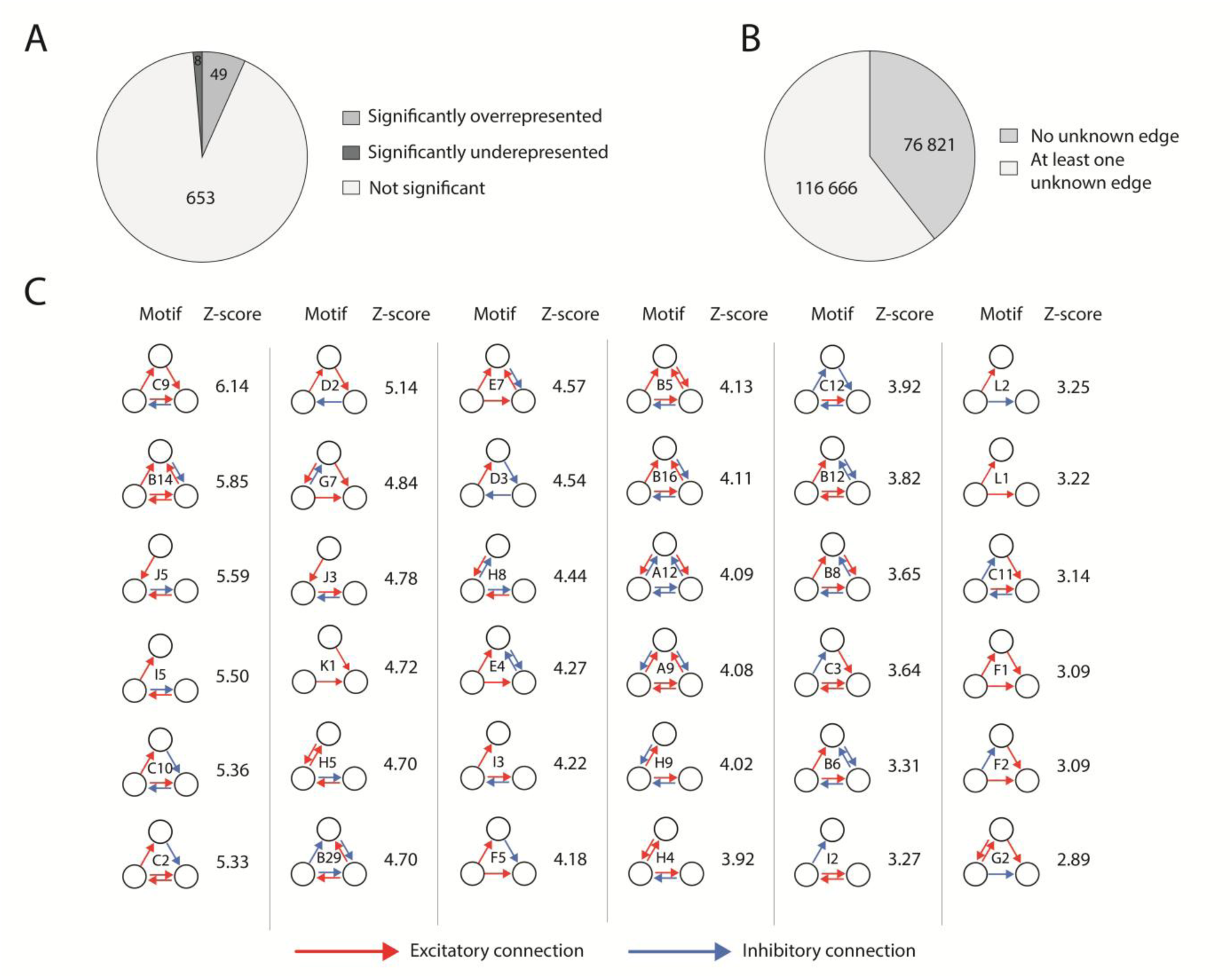
Results of partial motif analysis. A) Distribution of all 710 possible signed three-node motifs. Over– and underrepresentation were assessed against 1,000 random networks using the Z-score as a quantitative measure. B) Distribution of partial motifs of the connectome according to the presence or absence of unknown edges. C) Significantly overrepresented partial motifs (without unknown edges) and their respective Z-scores. The red and blue arrows represent excitatory and inhibitory synapses, respectively. All data are available in Additional file 5.

### Alternative randomization method

When the null model generated by an alternative (edge polarity shuffling) randomization method (see Methods) was used, we observed a greater number of significantly overrepresented motifs than what was found with the primary (neurotransmitter shuffling) method (Additional file 6). A total of 147 motifs were significantly overrepresented, and 148 motifs were significantly underrepresented. We found that 70 of the overrepresented motifs had no unknown edges. Motif H5 was only significant when the neurotransmitter-shuffling randomization method was used. In contrast, only the alternative randomization revealed 55 motifs (without unknown edges) that were significantly over– or underrepresented in the connectome (Fig. 6A).

**Fig. 6.**
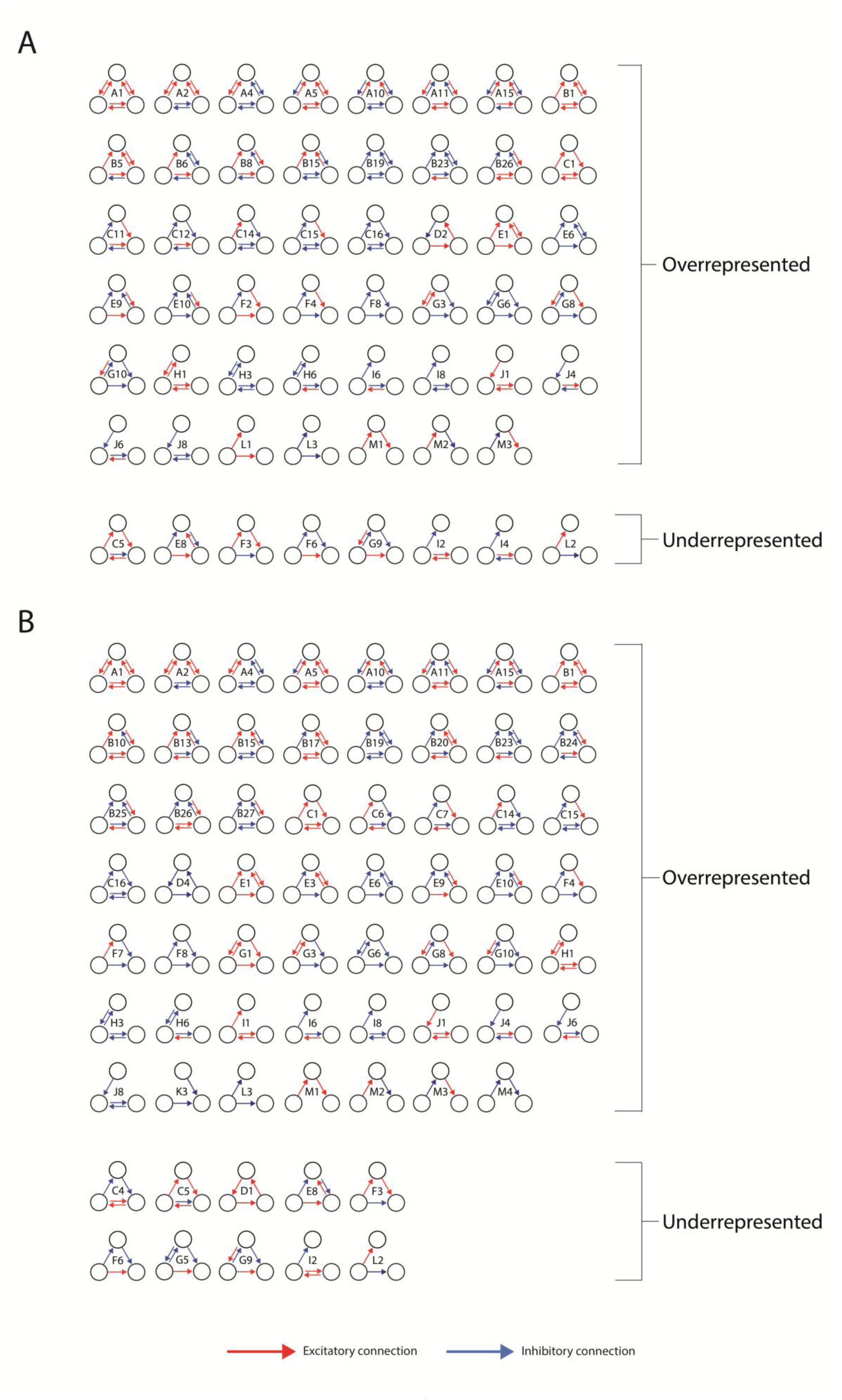
Induced and partial motif analysis using the alternative (edge polarity-shuffling) null model (n = 1000 random networks). A) Induced motifs (without unknown edges) identified to be significantly over– or underrepresented in the connectome solely by the alternative (edge polarity-shuffling) null model. With one exception (motif H5), all induced motifs that were significant when the neurotransmitter-shuffling method was used were also significant using this alternative null model. The red and blue arrows represent excitatory and inhibitory synapses, respectively. B) Partial motifs (without unknown edges) identified to be significantly over– or underrepresented in the connectome solely by the alternative null model. With one exception (motif G2), all partial motifs that were shown to be significant using the neurotransmitter-shuffling method were also significant using this alternative (edge polarity-shuffling) null model. The red and blue arrows represent excitatory and inhibitory synapses, respectively. All data are available in Additional File 6.

Using the alternative (edge polarity-shuffling) null model to address partial motif counts we observed an increased number of overrepresented motifs (Additional file 6). In this case, 179 motifs were significantly overrepresented, whereas 194 motifs were significantly underrepresented in the connectome. Among those, 88 and 10 motifs had no unknown edges, respectively. All but one (i.e., G2) motifs that were overrepresented using the neurotransmitter-shuffling randomization method also had a significant p-value while using the alternative null model as a comparison (Fig. 6B). In contrast to the primary model, using the alternative (edge polarity-shuffling) null model identified 10 significantly underrepresented motifs. Surprisingly, two of those motifs (motifs I2 and L2) were significantly overrepresented in the primary model.

### Validation

We performed perturbations on the network to evaluate the effect of major limitations on our results. We found that eliminating unknown connections (Additional file 9) did not significantly reduce the number of 18 induced motifs among the 23 that were previously identified as overrepresented in the connectome (78%). Similarly, during partial motif analysis, 27 of the 36 partial motifs previously found to be overrepresented did not have a significantly lower occurrence after the perturbation (75%).

To address potential errors in the predicted polarity data from our previous work [19], we generated another null model by randomly changing 10% of the edge signs in the network (Additional file 10. Doing that, we found that this process does not significantly lower the occurrence of more than 80% of the motifs reported to be overrepresented in the connectome (20 of the 24 induced motifs, 29 of the 36 partial motifs). However, the number of the coherent positive feed-forward motif (labelled as F1) significantly decreased during both induced and partial motif analysis.

### Feedforward and feedback loops

We analyzed feedforward and feedback motifs in detail, as they are known to be important in signaling networks (Table 1, references [25–32]). Since partial feedforward and feedback loops embedded in more complex motifs can function on their own, we used the results of partial motif counting for this analysis. We distinguished eight types of feedforward and four types of feedback loops, based on previously published concepts [6, 30], and then compared their occurrences in the connectome (Fig. 7A).

**Fig. 7.**
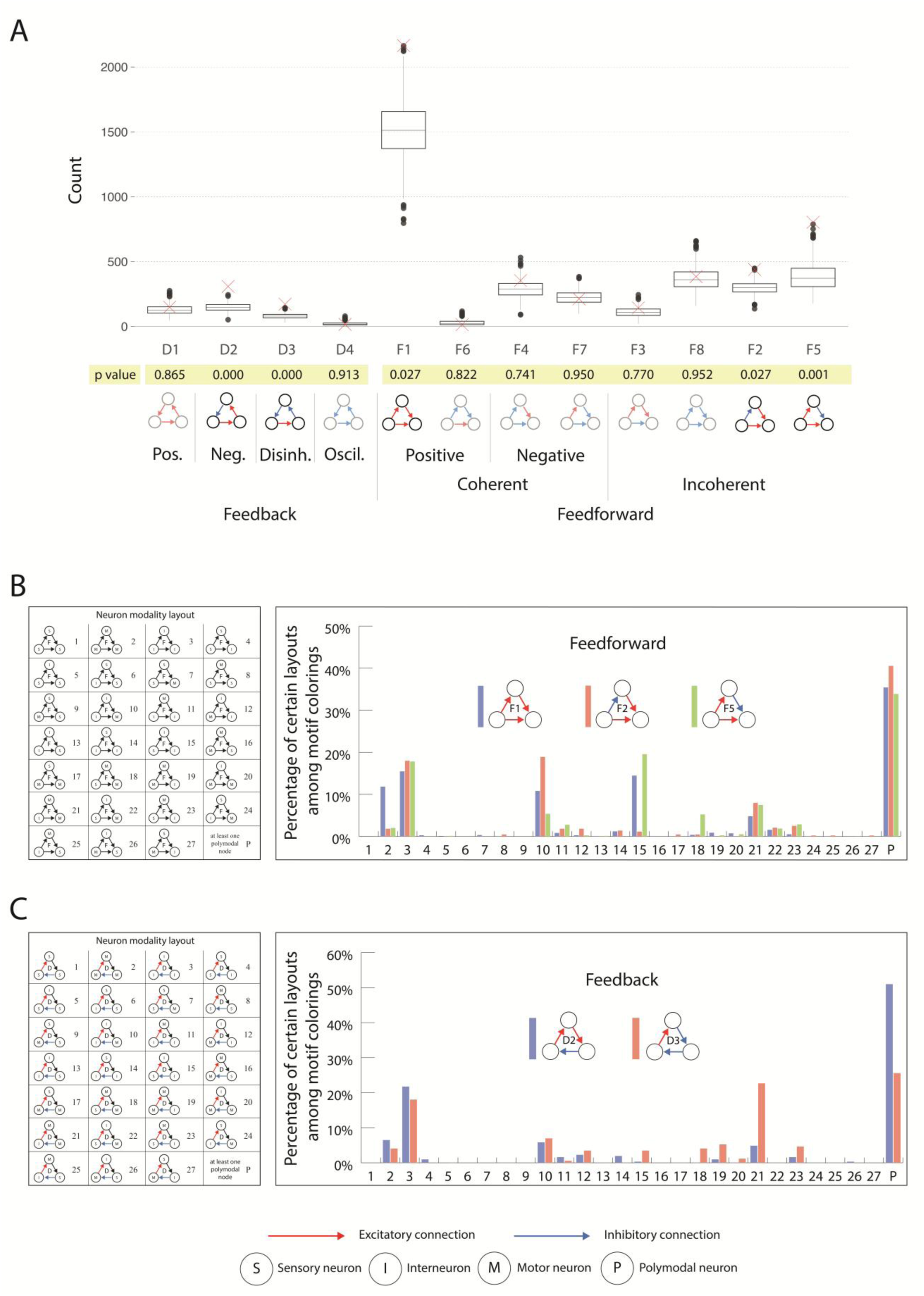
Feedforward and feedback motifs. A) Feedforward and feedback partial motif counts in the connectome (red X) compared with the null model of random networks (n = 1000, boxplot). Positive feedback (Pos.), negative feedback (Neg.), disinhibitory feedback (Disinh.), oscillatory feedback (Oscil.), positive feedforward, negative feedforward, and incoherent feedforward loops are shown. All data are available in Additional file 5. B) Distribution of significant partial feedback and feedforward motifs by neuron modality layouts. The neuron modalities are abbreviated as S (sensory neuron), I (interneuron), and M (motor neuron). Layouts containing at least one polymodal neuron (i.e., a neuron without one specific modality) were combined and labeled P. C) Distribution of significant partial feedback motifs by neuron modality layouts. The neuron modalities are abbreviated as S (sensory neuron), I (interneuron), and M (motor neuron). Layouts containing at least one polymodal neuron (i.e., a neuron without one specific modality) were combined and labeled P. Uncolored feedback motifs have rotational symmetry; however, colored motifs do not. Therefore, the red and blue arrows (corresponding to excitatory and inhibitory connections, respectively) of the neuron modality layout table define the exact rotational layouts of the motifs.

**Table 1.** Feedback and feedforward signed motifs. Table 1. Types and roles of signed feedforward and feedback motifs in biological networks. Motifs in **bold** are significantly overrepresented in the *C. elegans* connectome.

|  | Motif | Type | Function | References |
| --- | --- | --- | --- | --- |
| Feedback | D1 | Positive | Amplification, information processing | [25] |
|  | D2 | Negative | Oscillator, stabilizing | [25] |
|  | D3 | Disinhibitory<br>(positive) | Amplification, creating a dense response | [26, 27] |
|  | D4 | Oscillatory<br>(negative) | Biological repressilator, oscillator | [25, 28] |
| Feedforward | F1 | Coherent<br>positive | Sign-sensitive delay/persistence detector / learning; overrepresented in many systems | [29, 30] |
|  | F2 | Incoherent | Pulse generator, sign-sensitive accelerator. | [30] |
|  | F3 | Incoherent | Pulse generator, sign-sensitive accelerator. | [30] |
|  | F4 | Coherent<br>negative | Sign-sensitive delay | [30] |
|  | F5 | Incoherent | Pulse generator, response accelerator, fold-change detection in sensory systems, input normalization; overrepresented in many systems | [30, 31] |
|  | F6 | Coherent<br>disinhibitory | Sign-sensitive delay, epileptic seizures | [30, 32] |
|  | F7 | Coherent<br>negative | Sign-sensitive delay | [30] |
|  | F8 | Incoherent | Pulse generator, sign-sensitive accelerator | [30] |

Addressing feedforward loops (Fig. 7A), we found that the F1 coherent positive feedforward loop and the F2 and F5 incoherent feedforward loops were overrepresented (Z-scores of 3.09, 3.09, and 4.18, respectively). Moreover, the D2 negative feedback (Z-score = 5.14) and the D3 feedback disinhibition (Z-score = 4.54) loops were also highly overrepresented. Positive feedback (D1) and coherent negative feedforward (F4, F7) loops were not overrepresented.

In the next step, we further analyzed the distributions of neuron modalities (i.e., sensory, motor, inter, polymodal) among the significant feedforward and feedback loops (the list of neuron modalities is provided in Additional file 7). The overrepresented feedforward motifs occur at a high, yet similar, level among interneurons (layout #3). Excluding that, we found a pattern in which specific modality layouts (excluding polymodal-containing motifs) dominated each feedforward motif (Fig. 7B). For the F2 (incoherent) motif, the dominant modality layout was #10, which represents inter>inter inhibition and inter>motor excitation patterns. For the F5 (also incoherent) motif, the dominant layout was also #15, which in the case of this motif is a sensory>inter excitatory and inter>inter inhibitory layout.

In the case of feedback loops (Fig. 7C), both D2 negative feedback and D3 disinhibitory loops were frequently observed in the interneuron-only layout (#3). On the other hand, the D3 feedback loop was dominant in modality layout #21 (representing inter>motor excitation, motor->motor inhibition, and motor>inter inhibition), whereas D2 was not. Surprisingly, we did not observe a single instance of the archetypal three-layer negative feedback loop (i.e., layout #22: sensory > inter and inter > motor excitation, and motor > sensory inhibition). Modality layouts with at least one polymodal neuron occurred more frequently across all motifs, but were presented as a single layout to simplify interpretation.

### Two-node colored motif analysis considering neuronal modalities

We performed induced and partial motif analysis of two-node motifs, while distinguishing neuron modalities, as was done with three-node feedforward and feedback loops (Additional file 8). During induced motif analysis, we observed 15 significantly over– and 8 underrepresented motifs. Ten of the overrepresented and 2 of the underrepresented motifs had no unknown edges (Fig. 8). During partial motif enumeration, we identified 19 significantly overrepresented and 12 underrepresented motifs, 13 and 4 of which had only excitatory and inhibitory edges, respectively (Fig. 8). All the motifs found to be significant in the induced motif analysis were also significant in the partial motif analysis.

**Fig. 8.**
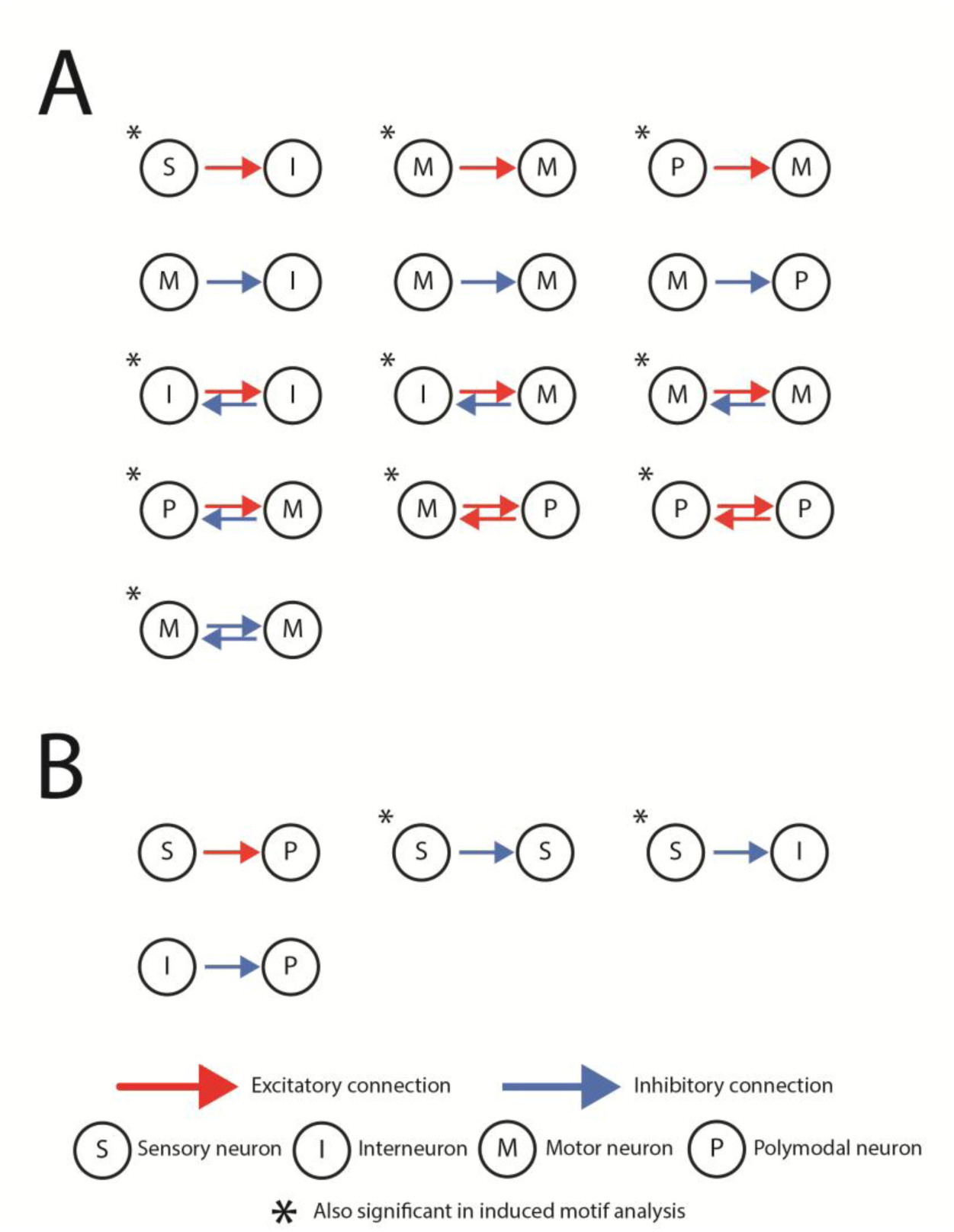
Two-node motif analysis with neuron modalities, using the neurotransmitter-shuffling null model (n = 1000). Motifs marked with an asterisk were also significantly overrepresented in the induced motif analysis. The neuron modalities are abbreviated as S (sensory neuron), I (interneuron), M (motor neuron), and P (polymodal neuron). The red and blue arrows represent excitatory and inhibitory synapses, respectively. A) Significantly overrepresented two-node partial motifs differentiating neuron modalities. B) Significantly underrepresented two-node partial motifs differentiating neuron modalities. All data are available in Additional file 8.

## Discussion

The neuronal network of *C. elegans* is the first fully reconstructed connectome [9], enabling the detailed analysis of structural characteristics, such as the abundance of network motifs in the nervous system. We conducted, for the first time, edge-polarity based signed motif analysis using the previously published signed connectome of *C. elegans*.

As we expected, partial motif analysis revealed some overrepresented motifs that were not found to be significant during induced motif enumeration. This is consistent with previous studies showing that simple motifs (such as feed-forward and feedback loops) are often embedded in complex structures [33, 34], including mixed feed-forward loops and complete graphs.

We found that the previously reported high number of feedforward loops [34–36] is disproportionately distributed among the different edge sign patterns. Compared with random networks, two types of incoherent feedforward loops are overrepresented. Incoherent feedforward loops are overrepresented in many systems [3, 37–39] and function as sign-sensitive accelerators that play key roles in network information processing [30]. In the *C. elegans* connectome, they are found in circuits responsible for chemotaxis [40]. The negative feedback loop is also well known in various networks [41, 42]. In the *C. elegans* connectome, negative feedback loops have been observed in mechanosensory processing [43], chemotaxis [44, 45], and (along with positive feedback loops) rhythmic forward locomotion [46]. We also found an overrepresentation of the disinhibitory feedback motif, which has a more complex function: it helps in balancing excitation and inhibition in a signed network. This can lead to novel response amplification and a dense population response [27]. Because it has two distinct steady states [6], a disinhibitory feedback motif can act as a biological switch [47]. We identified 172 three-neuron circuits in the connectome that form disinhibitory feedback loops. Experimental activation or inhibition of one neuron within these circuits should produce coordinated changes in the activity of the other two neurons and may induce transitions between alternative stable circuit states.

While testing the effect of perturbations in the polarity database, we observed a significant decrease in the number of the coherent positive feedforward motif (F1). Therefore, the overrepresentation of this motif may be due to any potential error in the predicted polarity data, and we should refrain from treating it as a core organizational feature of the *C. elegans* connectome. However, coherent positive feedforward loops have been reported in the intracellular adaptation to thermosensation [48] and in the egg-laying circuit [49] of the animal. For the F1 coherent positive feedforward loop, the most common modality layouts were #2 and #15; the former represents the feedforward loop of motor neurons, and the latter denotes feedforward processing from sensory neurons to interneurons. Therefore, the functional importance of this motif can be experimentally verified by activating sensory neurons, recording interneuron activity, and observing the feedforward processing in circuits described as coherent feedforward motifs.

Labelling 20% of the neurons as polymodal resulted in many motifs containing at least one polymodal node. However, we observed a distinct neuron modality layout for each overrepresented feedback loop (Fig. 7). Surprisingly, we did not find a single occurrence of the canonical negative feedback loop (numbered as layout #22: sensory>inter, inter>motor, and sensory>motor connections). The classical downstream feedforward signaling was also not found to be overrepresented. The lack of this type of feedforward loops has been previously shown in the connectome [35, 50]. The lack of these motifs is partly because many neurons are responsible for multiple different behaviors [51], hence, forming direct sensory-inter-motor loops is unlikely. Instead, our findings suggest that while the canonical negative feedback exists globally in the network [19], it is more likely to be formed by indirect connections between neurons, which routes could be revealed by circuit mapping or perturbation experiments. Meanwhile, information integration occurs at the level of interneurons via feedforward loops (Fig. 7). This is consistent with previous studies describing strong information integration within the nerve ring. [52–54].

Even though the structure of motifs F2 and F5 looks similar (incoherent feedforward loops with an excitatory direct pathway), their kinetics are different [30]. Therefore, we could also identify different characteristic layouts for each (layout #10 and layout #15, respectively). We observed that the F2 motif is crucial for motor neuron activation, whereas F5 contributes to fold-change sensory detection in interneurons. While these specific phenomena are yet to be observed experimentally in the *C. elegans* connectome, our results correlate with findings in other nervous systems [55, 56]. Even though the occurrence of incoherent feedforward partial motif F5 significantly reduced by resolving unknown connections (Additional file 9), the resulting frequency still indicated significant overrepresentation; thus, the motif did not fail the validity test. However, the true functional relevance of motif F5 can be verified by potential electrophysiological experiments, such as observing the fold-change sensory detection of interneurons by activating the sensory neurons of these circuits, then measuring interneuron activity levels.

Of the two randomization methods, the primary (neurotransmitter-shuffling) randomization method yielded a biologically meaningful null model that hypothesized that polarity assignment in the connectome is independent of its structure. Our findings suggest that polarity distribution and assignment in the *C. elegans* connectome are structurally non-random. Our alternative, edge polarity shuffling randomization method yielded an excessive number of significant motifs in the connectome. However, all significant motifs of the neurotransmitter-shuffling randomization method – except motif H5 in induced motif analysis, and G2 in partial motif analysis – were also significant in the edge polarity shuffling method. This enhances the robustness and validity of our findings. It is worth noting that the connectivity structure of the connectome varies among individual animals [54] and changes during development [57]. Therefore, additional methods that randomize connections could provide further insight into the importance of network motifs. Furthermore, the exact biological function of each over-represented signed motif should be experimentally assessed.

We found that 159 motifs were absent from the connectome. However, since these motifs also had low occurrence in the null model, they are not considered to be significantly underrepresented in the network. A total of 150 of the 159 missing motifs have at least 5 edges. Unsurprisingly, the occurrence of more complex motifs is relatively low in both induced and partial analyses. Therefore, only motifs with a biological function should be represented in the network. We found that more than 80% of missing complex motifs without unknown edges have an excitatory/inhibitory ratio <1, which contradicts our understanding of sign balance both generally in biological networks and the *C. elegans* connectome [19]. Data limitations also led to the loss of motifs. In addition to unknown edges, resolving edges with complex predicted polarity also influenced the motif landscape. As partial motifs can be embedded in subgraphs with more connections, excitation and inhibition can similarly occur through synapses labelled as complex in our previous work [19].

Previous studies [34, 35] have shown the overrepresentation of symmetrical three-node motifs with a bidirectional edge (labeled in our work as motifs E and G). These motifs are often labeled as “mixed feedforward loops” because two partial feedforward loops are embedded in their structure. We found that only 4 of the 20 possible colorings of these motifs were significantly overrepresented (Fig. 5C). Surprisingly, even though the structures of motifs E and G are symmetrical, 3 of the 4 overrepresented motifs are asymmetrical in color (Table 2). Breaking down these mixed feedforward loops to their elementary units (partial motifs), we find that two of them have functionally different components (Table 2).

**Table 2.**
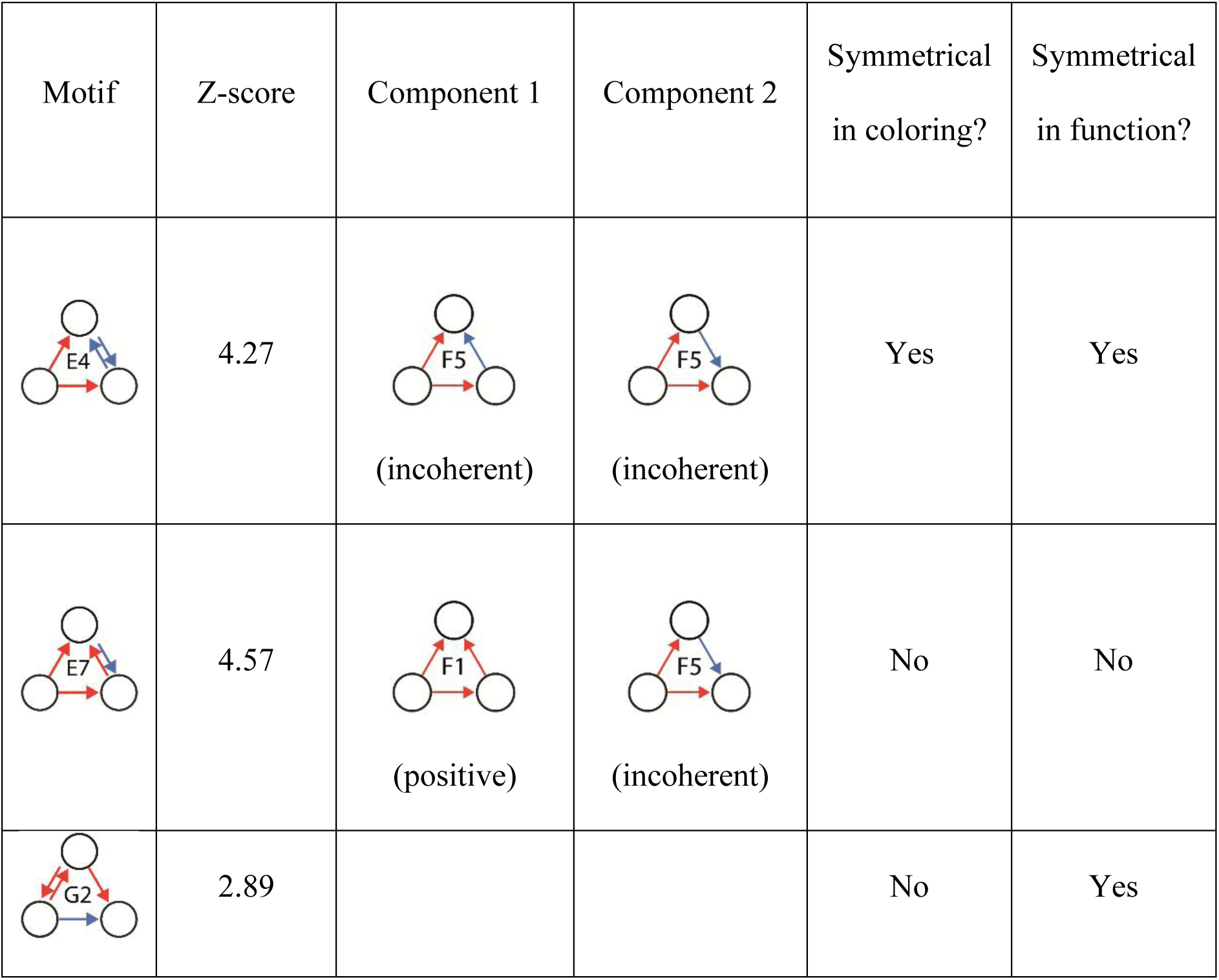
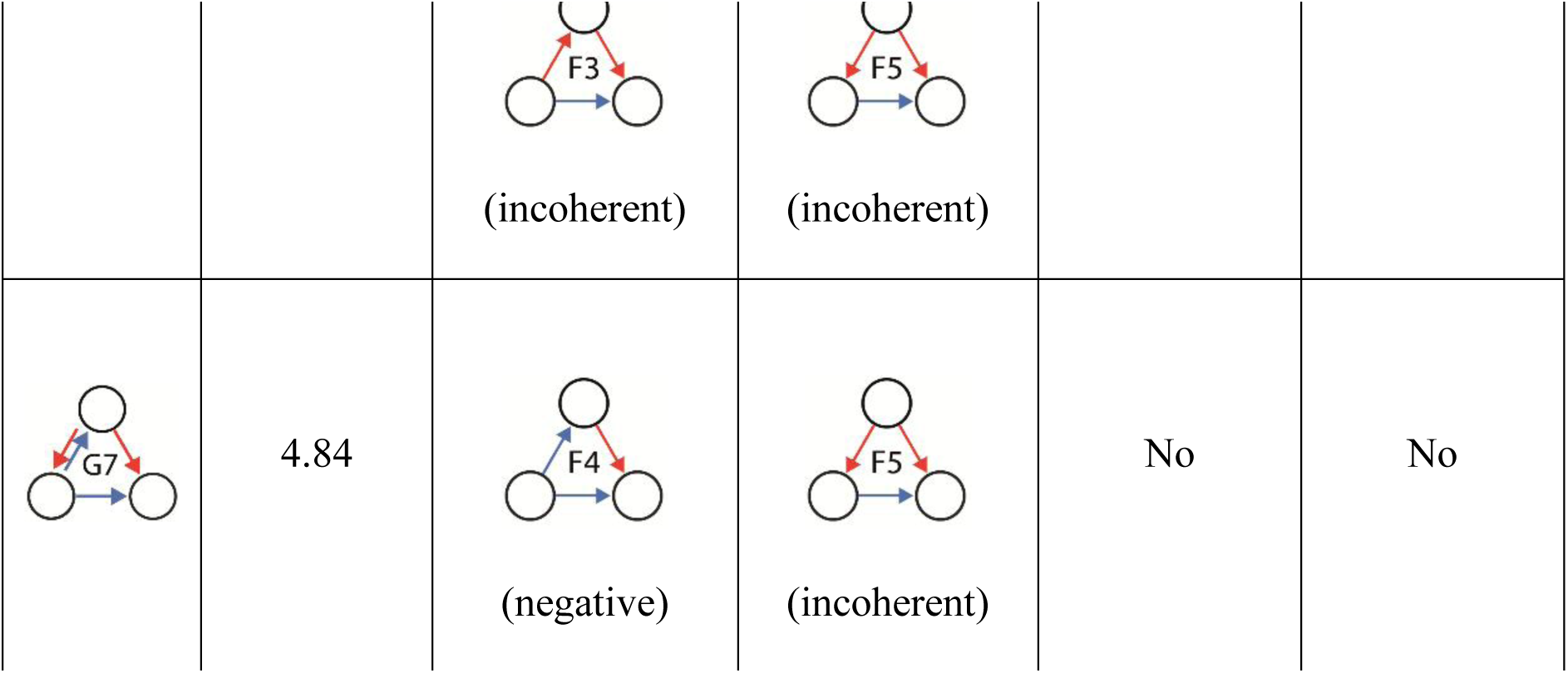
Components of significantly overrepresented mixed feedforward loops. Table 2. Despite the animal’s bilateral symmetry, most of the overrepresented mixed feedforward motifs exhibit asymmetrical coloring. Only half of the overrepresented mixed feedforward motifs contain partial feedforward loops with identical function

| Motif | Z-score | Component 1 | Component 2 | Symmetrical in coloring? | Symmetrical in function? |
| --- | --- | --- | --- | --- | --- |
|  | 4.27 | (incoherent) | (incoherent) | Yes | Yes |
|  | 4.57 | (positive) | (incoherent) | No | No |
|  | 2.89 |  |  | No | Yes |
|  |  | (incoherent) | (incoherent) |  |  |
|  | 4.84 | (negative) | (incoherent) | No | No |

Dissolving mixed feedforward loops into their components provides insight into their roles. These motifs are made of two partial feedforward components. Even though the structures of these motifs are symmetrical, two of the motifs (motifs E7 and G7) are built by feedforward loops with different modalities. Three of the 4 overrepresented mixed loops are asymmetrical in color (motifs E7, G2, and G7). The red and blue arrows represent excitatory and inhibitory synapses, respectively.

In brain networks, motifs with bidirectional connections can induce zero-lag synchrony in neuronal spiking models [58]. While 28 of the 36 overrepresented motifs contained at least one bidirectional edge (Fig. 5C), 13 motifs had more than one. Unfortunately, modeling bidirectional connections is quite challenging [59] because of the complex temporal aspects of neurotransmission.

While three-node motifs are the standards for motif analysis [3], two-node motifs can act as a smaller, more direct level of connectivity [60]. Additionally, simultaneously signing the nodes (based on the neuronal modality) and the edges (based on the synaptic polarity) can provide novel insight into network dynamics. Using this method, we demonstrated several unique functional characteristics of the connectome, such as downstream (sensory-to-interneuron) activation and motor-to-interneuron inhibition. These mechanisms were previously described in the *C. elegans* connectome [61, 62]. Motor-to-motor neuron inhibition is possibly related to contralateral motor neuron inhibition, which is necessary for physiological locomotion. On the other hand, forward activation and backward inhibition between two motor neurons might occur between ipsilateral motor neurons. Although two-node motifs have marginal importance in network dynamics, a similar analysis of edge– and node-signed three-node motifs would reveal further functional information. However, extending this method to three-node motifs would have produced an excessively large number of motifs, making interpretation difficult.

Our study has several limitations. Since we solely used the connectome described in the WormWiring connectome reconstruction (http://wormwiring.org), our reported motif landscape may differ from what could be observed in different individual animals [54] or during development [57]. Due to limited data on synaptic polarity, only 40% of all three-node motifs in the connectome (graphlets with edges that are either excitatory or inhibitory) were suitable for interpretation. While resolving these connections did not significantly lower the ratio of most of the motifs that were reported to be overrepresented, a potential future curation and completion of polarity data could substantially improve the analysis of the motif landscape. New connectivity data are still emerging [52, 57], and recent experimental results on synaptic polarity and gene expression have also been reported [63]. The high number of complex connections [19] also implies that our results can be complemented once the role of those synapses is better understood. Resolving complex connections to excitatory and inhibitory edges certainly affected our results. However, the exact function of complex synapses in motifs remains to be determined, while treating them as an additional edge color would result in an almost fourfold increase in the number of distinct motif colorings.

Focusing on three-node motifs is another limitation of our work. However, since the number of possible four-node signed motifs is extremely high (more than 700,000 motifs), it would be difficult to interpret the results of four-node motif enumeration. Furthermore, there are only a handful of four-node motifs with a defined biological function. Therefore, similarly to previous studies [10, 13, 18], we focused on three-node motifs in the connectome.

Focusing solely on the chemical connections by excluding electrical synapses is also a limitation of our study. These connections can be considered as bidirectional edges in the network [64], but rather than having polarity, they are usually treated as sign-symmetric electrical couplings whose effect emerges from surrounding network properties [65]. Therefore, even though electrical synapses can be integrated into motif analysis of the unsigned network [10, 15, 50], their nature did not fit into our definition of signed motifs. However, a planned future project will extend this work by analysing feedforward and feedback loops in the connectome by the presence of potential electrical synapses between their nodes. While the number of electrical connections is relatively low in these structures, they are disproportionately distributed among the different colored motifs. Since the biological reason behind this is yet to be understood, we aim to explore this aspect of motif analysis in the future.

Finally, our work does not include the extrasynaptic neuropeptide and monoamine systems, which can also mediate feedback control in the network [66–70]. Since these molecules are transported without an anatomical structure, it is impossible to illustrate their connectivity pattern as a hard-wired network [68]. The way these molecules function is governed by complex regulatory mechanisms [69, 70]. Although monoamines and neuropeptides certainly play a role in the regulation of the connectome [71], we could not assume that these connections have the same influence on neuronal motif structures as chemical synapses.

Despite our limitations, our work provides novel insight into the motif landscape of the *C. elegans* nervous system. Compared with previous studies (Table 3; references [10, 13, 16–18]), our model uses a concept that assigns signs to the edges of the connectome based on their roles in synaptic transmission. Edge-signed motif analyses have been attempted before (Table 3), but since neurotransmitters do not determine synaptic polarity on their own [19], our work offers a more functional approach to motif analysis.

**Table 3.**
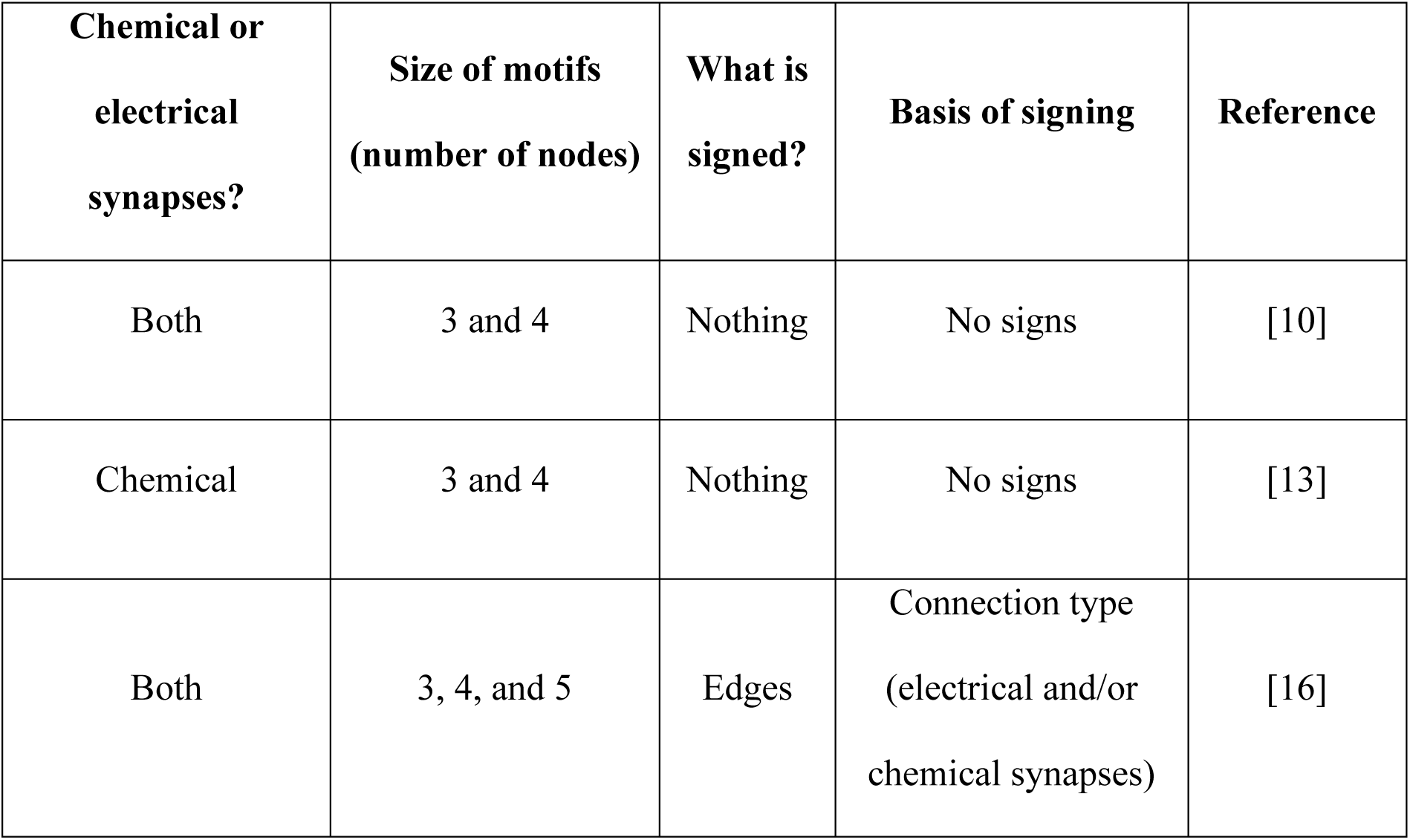
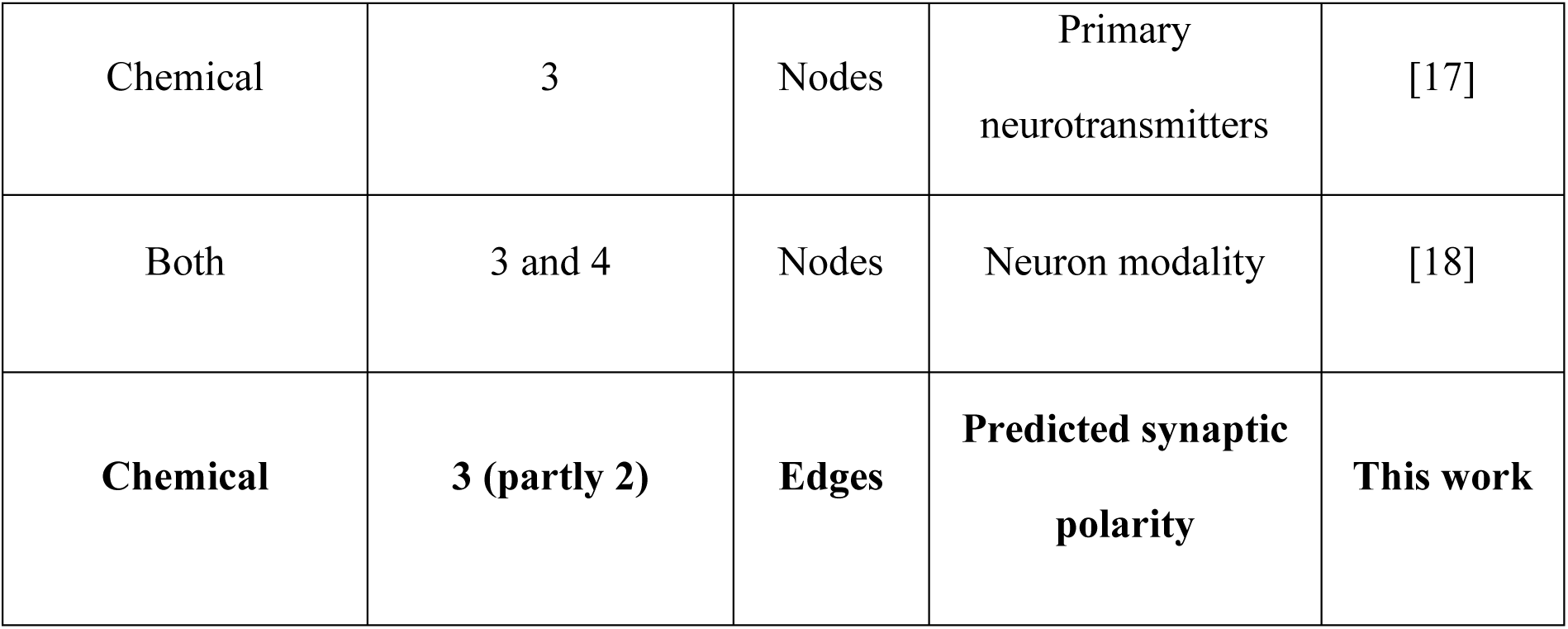
Comparison of motif studies in the *C. elegans* connectome. Table 3. There were multiple motif studies on the *C. elegans* connectome. Some of them were signed analyses that labeled the network’s nodes or edges. However, our study is the first to use synaptic polarities for signed motif analysis.

### Conclusions

Although a form of neurotransmitter-based motif analysis has been explored before [17], to our knowledge, our study is the first to apply edge-polarity-based signed motif analysis to a fully mapped connectome. Using predicted edge-polarities, our work revealed insights into the functional architecture of the *C. elegans* neuronal network. The overrepresentation of certain motif types, particularly incoherent feedforward and negative feedback loops, suggests that these motifs may contribute significantly to the network’s information processing and homeostatic functions, although their specific biological relevance remains to be experimentally determined. Our results correlate with experimental data on motif functions, such as incoherent feedforward and negative feedback motifs. To validate our results, the performed sensitivity analyses suggest that overrepresented motif structures remain stable after perturbation. Identifying these circuits in the network can provide potential targets for further experimental validation of our findings. Future work that incorporates emerging synaptic polarity data and expands beyond three-node motifs could yield even deeper insights into the organizational principles of neural networks. As motif analysis techniques evolve and are applied to larger-scale connectomes, our findings may offer a foundation for exploring how distinct motif configurations support the complex functions of diverse nervous systems.

## Declarations

## Acknowledgements

The authors thank Istvan Kovacs (Northwestern University, Department of Physics and Astronomy) and Balazs Hangya (HUN-REN Institute of Experimental Medicine) for their valuable discussions, and all members of the LINK group for their input.

## Consent for Publication

### Funding

This work was also supported by the Thematic Excellence Programme (Tématerületi Kiválósági Program, TKP2021-EGA-24) of the Ministry for Innovation and Technology in Hungary, within the framework of the Molecular Biology thematic program at Semmelweis University.

### Data availability

All data generated or analysed during this study are included in this published article, its supplementary information files and publicly available repositories [21].

### Authors’ Contributions

G.S.S. contributed to the study conception, design, development of the algorithm, analysis, and writing of the manuscript; A.G. contributed to the study conception and analysis; Z.V. contributed to the interpretation of the data; P.C. contributed to the study conception and interpretation of the data and contributed to the manuscript; B.G.F. contributed to study conception, data analysis, and manuscript writing. All authors read and approved the final manuscript

### Ethics Approval

Non-applicable for the current studies

### Consent to Participate

Non-applicable for the current studies

## Description of supplementary data

**Additional file 1**

**File format: Excel spreadsheet (.xlsx)**

**Title of data: Polarity of edges in the connectome and random networks**

**Description of data: The edge polarities of the connectome (labeled “Pred”) were obtained from our previous work** [19]**. Edge polarity was also predicted in the 1,000 random networks of the null model.**

**Additional file 2**

**File format: Excel spreadsheet (.xlsx)**

**Title of data: Motif codes and dictionary**

**Description of data: All possible edge codes (as described in the Methods section) correspond to a single motif (see the ‘Motif definitions’ sheet). As isomorph motifs should be labeled the same, the ‘Unique motifs’ sheet contains all the different motifs and all their possible edge layouts. The ‘Partial Motifs’ list lists all the partial motifs for every induced motif.**

**Additional file 3**

**File format: Excel spreadsheet (.xlsx)**

**Title of data: Motifs in the connectome**

**Description of data Using the motif dictionary and the partial motif list from Additional file 2, the induced and partial motifs of the connectome were enumerated as described in Additional file 1.**

**Additional file 4**

**File format: Excel spreadsheet (.xlsx)**

**Title of data: Induced motif counts and Z-scores in the connectome and null models Description of data: Using the edge lists from Additional file 1 and the motif dictionary described in Additional file 2, our induced motif enumerating tool counted the occurrences of all 710 different 3-node-colored motifs in the connectome and in the 1,000 random networks.**

**Additional file 5**

**File format: Excel spreadsheet (.xlsx)**

**Title of data: Partial motif counts and Z-scores in the connectome and null models Description of data: Using the edge lists from Additional file 1 and the motif dictionary described in Additional file 2, our partial motif enumerating tool was able to count the occurrence of all 710 different 3-node colored motifs in the connectome and 1,000 randomly generated networks.**

**Additional file 6**

**File format: Excel spreadsheet (.xlsx)**

**Title of data: Motif analysis of the connectome via the alternative null model Description of data: Using the edge polarity data from the alternative randomization method (see the ‘Edges’ sheet), we were able to perform induced and partial motif counting. The process was the same as that described in Additional files 4 and 5, except that the edge polarity data were different.**

**Additional file 7**

**File format: Excel spreadsheet (.xlsx)**

**Title of data: Neuron modalities**

**Description of data: List of neuron modalities based on the data from** http://wormatlas.org

**Additional file 8**

**File format: Excel spreadsheet (.xlsx)**

**Title of data: Two-node motif analysis of the connectome considering neuronal modalities Description of data: Using the edge lists from Additional file 1, the two-node motif dictionary and neuron modality tables described here, our motif enumerating tool was able to count the occurrence of all 126 different two-node motifs in the connectome and 1,000 randomly generated networks.**

**Additional file 9**

**File format: Excel spreadsheet (.xlsx)**

**Title of data: Validation of our results by resolving unknown connections**

**Description of data Using the edge polarity data from our previous work, we randomly assigned a polarity to unknown connections while maintaining the excitatory:inhibitory ratio of the network. After generating 1,000 networks, we observed the changes in both induced and partial motif occurrences to evaluate the stability of our results.**

**Additional file 10**

**File format: Excel spreadsheet (.xlsx)**

**Title of data: Validation of our results by randomizing 10% of edge polarities in the network**

**Description of data: Using the data from our previous work, we randomly altered 10% of the edge polarities. After generating 1,000 networks, we observed the changes in both induced and partial motif counts to evaluate the stability of our results.**

## References

1. Bassett DS, Bullmore ET. Small-world brain networks revisited. Neuroscientist. 2017;23:499–516. 10.1177/1073858416667720.

2. Bullmore E, Sporns O. Complex brain networks: graph theoretical analysis of structural and functional systems. Nat Rev Neurosci. 2009;10:186–98. 10.1038/nrn2575.

3. Milo R, Shen-Orr S, Itzkovitz S, Kashtan N, Chklovskii D, Alon U. Network motifs: Simple building blocks of complex networks. Science (80-). 2002;298:824–7. 10.1126/science.298.5594.824.

4. Sporns O, Kötter R. Motifs in brain networks. PLoS Biol. 2004;2:e369. 10.1371/journal.pbio.0020369.

5. McDonnell MD, Yaveroğlu ÖN, Schmerl BA, Iannella N, Ward LM. Motif-role-fingerprints: The building-blocks of motifs, clustering-coefficients and transitivities in directed networks. PLoS One. 2014;9:e114503. 10.1371/journal.pone.0114503.

6. Alon U. Network motifs: theory and experimental approaches. Nat Rev Genet. 2007;8:450–61. 10.1038/nrg2102.

7. Ryan K, Lu Z, Meinertzhagen IA. The CNS connectome of a tadpole larva of *Ciona intestinalis* (L.) highlights sidedness in the brain of a chordate sibling. Elife. 2016;5:1–34. 10.7554/eLife.16962.

8. Schlegel P, Yin Y, Bates AS, Dorkenwald S, Eichler K, Brooks P, et al. Whole-brain annotation and multi-connectome cell typing of *Drosophila*. Nature. 2024;634:139–52. 10.1038/s41586-024-07686-5.

9. White JG, Southgate E, Thomson JN, Brenner S. The structure of the nervous system of the nematode *Caenorhabditis elegans*. Philos Trans R Soc London B, Biol Sci. 1986;314:1–340. 10.1098/rstb.1986.0056.

10. Varshney LR, Chen BL, Paniagua E, Hall DH, Chklovskii DB. Structural properties of the *Caenorhabditis elegans* neuronal network. PLoS Comput Biol. 2011;7:e1001066. 10.1371/journal.pcbi.1001066.

11. Itzkovitz S, Alon U. Subgraphs and network motifs in geometric networks. Phys Rev E. 2005;71:026117. 10.1103/PhysRevE.71.026117.

12. Song S, Sjöström PJ, Reigl M, Nelson S, Chklovskii DB. Highly nonrandom features of synaptic connectivity in local cortical circuits. PLoS Biol. 2005;3:e68. 10.1371/journal.pbio.0030068.

13. Liu J, Yuan Y, Zhao P, Liu G, Huo H, Li Z, et al. Change of motifs in *C. elegans* reveals developmental principle of neural network. Biochem Biophys Res Commun. 2022;624:112–9. 10.1016/j.bbrc.2022.07.108.

14. Sharma D, Renz M, Hövel P. Discovering motifs to fingerprint multi-layer networks: a case study on the connectome of *C. elegans*. Eur Phys J B. 2025;98. 10.1140/epjb/s10051-024-00848-4.

15. Liu J, Yuan Y, Zhao P, Gu X, Huo H, Li Z, et al. Neuronal motifs reveal backbone structure and influential neurons of neural network in *C. elegans*. J Complex Networks. 2023;11. 10.1093/comnet/cnad013.

16. Matejek B, Wei D, Chen T, Tsourakakis CE, Mitzenmacher M, Pfister H. Edge-colored directed subgraph enumeration on the connectome. Sci Rep. 2022;12:11349. 10.1038/s41598-022-15027-7.

17. Pereira L, Kratsios P, Serrano-Saiz E, Sheftel H, Mayo AE, Hall DH, et al. A cellular and regulatory map of the cholinergic nervous system of *C. elegans*. Elife. 2015;4. 10.7554/eLife.12432.

18. Qian J, Hintze A, Adami C. Colored motifs reveal computational building blocks in the *C. elegan*s Brain. PLoS One. 2011;6:e17013. 10.1371/journal.pone.0017013.

19. Fenyves BG, Szilágyi GS, Vassy Z, Sőti C, Csermely P. Synaptic polarity and sign-balance prediction using gene expression data in the *Caenorhabditis elegans* chemical synapse neuronal connectome network. PLOS Comput Biol. 2020;16:e1007974. 10.1371/journal.pcbi.1007974.

20. Hardege I, Morud J, Courtney A, Schafer WR. A novel and functionally diverse class of acetylcholine-gated ion channels. J Neurosci. 2023;43:1111–24. 10.1523/JNEUROSCI.1516-22.2022.

21. Szilágyi G. Colored motif analysis project of the *C. elegans* connectome. Zenodo. 10.5281/zenodo.19682792.

22. Váša F, Mišić B. Null models in network neuroscience. Nat Rev Neurosci. 2022;23 August:493–504. 10.1038/s41583-022-00601-9.

23. Schbath S, Lacroix V, Sagot M-F. Assessing the exceptionality of coloured motifs in networks. EURASIP J Bioinforma Syst Biol. 2009;2009:1–9. 10.1155/2009/616234.

24. Xia F, Wei H, Yu S, Zhang D, Xu B. A survey of measures for network motifs. IEEE Access. 2019;7:106576–87. 10.1109/ACCESS.2019.2926752.

25. Cinquin O, Demongeot J. Positive and negative feedback: Striking a balance between necessary antagonists. J Theor Biol. 2002;216:229–41. 10.1006/jtbi.2002.2544.

26. Liu Z, Le Q, Lv Y, Chen X, Cui J, Zhou Y, et al. A distinct D1-MSN subpopulation down-regulates dopamine to promote negative emotional state. Cell Res. 2022;32:139–56. 10.1038/s41422-021-00588-5.

27. Schulz A, Miehl C, Berry MJ, Gjorgjieva J. The generation of cortical novelty responses through inhibitory plasticity. Elife. 2021;10. 10.7554/eLife.65309.

28. Elowitz MB, Leibler S. A synthetic oscillatory network of transcriptional regulators. Nature. 2000;403:335–8. 10.1038/35002125.

29. Keifer J, Houk JC. Modeling signal transduction in classical conditioning with network motifs. Front Mol Neurosci. 2011;4. 10.3389/fnmol.2011.00009.

30. Mangan S, Alon U. Structure and function of the feed-forward loop network motif. Proc Natl Acad Sci. 2003;100:11980–5. 10.1073/pnas.2133841100.

31. Goentoro L, Shoval O, Kirschner MW, Alon U. The incoherent feedforward loop can provide fold-change detection in gene regulation. Mol Cell. 2009;36:894–9. 10.1016/j.molcel.2009.11.018.

32. Birjandian Z, Narla C, Poulter MO. Gain control of γ frequency activation by a novel feed forward disinhibitory loop: Implications for normal and epileptic neural activity. Front Neural Circuits. 2013;7 NOV. 10.3389/fncir.2013.00183.

33. Gorochowski TE, Grierson CS, di Bernardo M. Organization of feed-forward loop motifs reveals architectural principles in natural and engineered networks. Sci Adv. 2018;4. 10.1126/sciadv.aap9751.

34. Azulay A, Itskovits E, Zaslaver A. The *C. elegans* connectome consists of homogenous circuits with defined functional roles. PLoS Comput Biol. 2016;12:1–16. 10.1371/journal.pcbi.1005021.

35. Reigl M, Alon U, Chklovskii DB. Search for computational modules in the *C. elegans* brain. BMC Biol. 2004;2:1–12. 10.1186/1741-7007-2-25.

36. Jarrell TA, Wang Y, Bloniarz AE, Brittin CA, Xu M, Thomson JN, et al. The connectome of a decision-making neural network. Science (80-). 2012;337:437–44. 10.1126/science.1221762.

37. Mank NN, Berghoff BA, Klug G. A mixed incoherent feed-forward loop contributes to the regulation of bacterial photosynthesis genes. RNA Biol. 2013;10:347–52. 10.4161/rna.23769.

38. Mangan S, Itzkovitz S, Zaslaver A, Alon U. The incoherent feed-forward loop accelerates the response-time of the gal system of *Escherichia coli*. J Mol Biol. 2006;356:1073–81. 10.1016/j.jmb.2005.12.003.

39. Shen-Orr SS, Milo R, Mangan S, Alon U. Network motifs in the transcriptional regulation network of *Escherichia coli*. Nat Genet. 2002;31:64–8. 10.1038/ng881.

40. Vidal-Saez MS, Vilarroya O, Garcia-Ojalvo J. A multiscale sensorimotor model of experience-dependent behavior in a minimal organism. Biophys J. 2024;123:1654–67. 10.1016/j.bpj.2024.05.008.

41. Krishna S. Structure and function of negative feedback loops at the interface of genetic and metabolic networks. Nucleic Acids Res. 2006;34:2455–62. 10.1093/nar/gkl140.

42. Yan Q, Zhu K, Zhang L, Fu Q, Chen Z, Liu S, et al. A negative feedback loop between JNK-associated leucine zipper protein and TGF-β1 regulates kidney fibrosis. Commun Biol. 2020;3. 10.1038/s42003-020-1008-z.

43. Kumar S, Sharma AK, Tran A, Liu M, Leifer AM. Inhibitory feedback from the motor circuit gates mechanosensory processing in *Caenorhabditis elegans*. PLoS Biol. 2023;21 9 September:1–20. 10.1371/journal.pbio.3002280.

44. Liu H, Wu JJ, Li R, Wang PZ, Huang JH, Xu Y, et al. Disexcitation in the ASH/RIM/ADL negative feedback circuit fine-tunes hyperosmotic sensation and avoidance in *Caenorhabditis elegans*. Front Mol Neurosci. 2023;16 March:1–18. 10.3389/fnmol.2023.1101628.

45. Larsch J, Flavell SW, Liu Q, Gordus A, Albrecht DR, Bargmann CI. A Circuit for gradient climbing in *C. elegans* chemotaxis. Cell Rep. 2015;12:1748–60. 10.1016/j.celrep.2015.08.032.

46. Zhao P, Wang B, Rong Y, Yuan Y, Liu J, Huo H, et al. The roles of feedback loops in the *Caenorhabditis elegans* rhythmic forward locomotion. PLOS Comput Biol. 2025;21 6 June:1–24. 10.1371/journal.pcbi.1013171.

47. Gardner TS, Cantor CR, Collins JJ. Construction of a genetic toggle switch in *Escherichia coli*. Nature. 2000;403:339–42. 10.1038/35002131.

48. Hill TJ, Sengupta P. Feedforward and feedback mechanisms cooperatively regulate rapid experience-dependent response adaptation in a single thermosensory neuron type. Proc Natl Acad Sci. 2024;121. 10.1073/pnas.2321430121.

49. Prashad S, Davison CA, Hammarlund M, Koelle MR. A positive feedback loop promotes the active internal state of the *C. elegans* egg-laying circuit. Curr Biol. 2025;35:5521–5533.e6. 10.1016/j.cub.2025.10.005.

50. Dvali S, Seguin C, Betzel R, Leifer AM. Diverging network architecture of the *C. elegans* connectome and signaling network. PRX Life. 2025;3:1–15. 10.1103/6wgv-b9m6.

51. Atanas AA, Kim J, Wang Z, Bueno E, Becker M, Kang D, et al. Brain-wide representations of behavior spanning multiple timescales and states in *C. elegans*. Cell. 2023;186:4134–4151.e31. 10.1016/j.cell.2023.07.035.

52. Emmons SW. Comprehensive analysis of the *C. elegans* connectome reveals novel circuits and functions of previously unstudied neurons. PLOS Biol. 2024;22:e3002939. 10.1371/journal.pbio.3002939.

53. Moyle MW, Barnes KM, Kuchroo M, Gonopolskiy A, Duncan LH, Sengupta T, et al. Structural and developmental principles of neuropil assembly in *C. elegans*. Nature. 2021;591:99–104. 10.1038/s41586-020-03169-5.

54. Brittin CA, Cook SJ, Hall DH, Emmons SW, Cohen N. A multi-scale brain map derived from whole-brain volumetric reconstructions. Nature. 2021;591:105–10. 10.1038/s41586-021-03284-x.

55. Wang H, Dewell RB, Zhu Y, Gabbiani F. Information in a collision detection circuit. 2019;28:1509–21. 10.1016/j.cub.2018.04.007.Feed-forward.

56. Borba C, Kourakis MJ, Schwennicke S, Brasnic L, Smith WC. Fold change detection in visual processing. Front Neural Circuits. 2021;15 August:1–15. 10.3389/fncir.2021.705161.

57. Witvliet D, Mulcahy B, Mitchell JK, Meirovitch Y, Berger DR, Wu Y, et al. Connectomes across development reveal principles of brain maturation. Nature. 2021;596:257–61. 10.1038/s41586-021-03778-8.

58. Gollo LL, Mirasso C, Sporns O, Breakspear M. Mechanisms of zero-lag synchronization in cortical motifs. PLoS Comput Biol. 2014;10:e1003548. 10.1371/journal.pcbi.1003548.

59. Park Y, Lee MJ, Son SW. Motif dynamics in signed directional complex networks. J Korean Phys Soc. 2021;78:535–41. 10.1007/s40042-021-00058-6.

60. Kim JR, Yoon Y, Cho KH. Coupled feedback loops form dynamic motifs of cellular networks. Biophys J. 2008;94:359–65. 10.1529/biophysj.107.105106.

61. Mclntire SL, Jorgensen E, Kaplan J, Horvitz HR. The GABAergic nervous system of *Caenorhabditis elegans*. Nature. 1993;364:337–41. 10.1038/364337a0.

62. Chalfie M, Sulston J, White J, Southgate E, Thomson J, Brenner S. The neural circuit for touch sensitivity in *Caenorhabditis elegans*. J Neurosci. 1985;5:956–64. 10.1523/JNEUROSCI.05-04-00956.1985.

63. Zhang G, Roberto NM, Lee D, Hahnel SR, Andersen EC. The impact of species-wide gene expression variation on *Caenorhabditis elegans* complex traits. Nat Commun. 2022;13:3462. 10.1038/s41467-022-31208-4.

64. Hall DH. Gap junctions in *C. elegans*: Their roles in behavior and development. Dev Neurobiol. 2017;77:587–96. 10.1002/dneu.22408.

65. Jin EJ, Park S, Lyu X, Jin Y. Gap junctions: Historical discoveries and new findings in the *Caenorhabditis elegans* nervous system. Biol Open. 2020;9:1–8. 10.1242/bio.053983.

66. Bentley B, Branicky R, Barnes CL, Chew YL, Yemini E, Bullmore ET, et al. The multilayer connectome of *Caenorhabditis elegans*. PLOS Comput Biol. 2016;12:e1005283. 10.1371/journal.pcbi.1005283.

67. Komuniecki R, Hapiak V, Harris G, Bamber B. Context-dependent modulation reconfigures interactive sensory-mediated microcircuits in *Caenorhabditis elegans*. Curr Opin Neurobiol. 2014;29:17–24. 10.1016/j.conb.2014.04.006.

68. Ripoll-Sánchez L, Watteyne J, Sun HS, Fernandez R, Taylor SR, Weinreb A, et al. The neuropeptidergic connectome of *C. elegans*. Neuron. 2023;111:3570–3589.e5. 10.1016/j.neuron.2023.09.043.

69. Rosikon KD, Bone MC, Lawal HO. Regulation and modulation of biogenic amine neurotransmission in *Drosophila* and *Caenorhabditis elegans*. Front Physiol. 2023;14 February:1–21. 10.3389/fphys.2023.970405.

70. Watteyne J, Chudinova A, Ripoll-Sánchez L, Schafer WR, Beets I. Neuropeptide signaling network of *Caenorhabditis elegans*: from structure to behavior. Genetics. 2024;228. 10.1093/genetics/iyae141.

71. Chalasani SH, Kato S, Albrecht DR, Nakagawa T, Abbott LF, Bargmann CI. Neuropeptide feedback modifies odor-evoked dynamics in *Caenorhabditis elegans* olfactory neurons. Nat Neurosci. 2010;13:615–21. 10.1038/nn.2526.

